# Exact analytical solution of the extensible freely jointed chain model

**DOI:** 10.1101/315051

**Authors:** Alessandro Fiasconaro, Fernando Falo

## Abstract

Based on classical statistical mechanics, we calculate analytically the length extension under a pulling force of a polymer modeled as a freely jointed chain with extensible bonds, the latter being considered as harmonic springs. We obtain an exact formula for the extension curve, as well as an independent high force approximation. These formulas can reproduce with high precision the experimental extension/force curves also at low values of the elastic constant of the spring, where previous proposals differ substantially. We successfully validate the analytical results together with the phenomenological expressions used in the literature by analyzing the precision of their fit on data obtained from Langevin simulations.

---

The stretching curve of a long polymer chain has been the subject of many theoretical and experimental studies. The first experiment was performed by the Bustamante group by using an optical tweezer to hang a single DNA molecule, with which they were able to stretch the filament under the application of a force [1]. Their results showed that the extension curve *vs* the applied force of a double stranded DNA (dsDNA), pulled at a small and intermediate force range, can be understood and depicted by means of the worm-like-chain model (WLC) [2], which consists in a semiflexible continuous beam. The WLC model can be discretized as a chain of beads connected by sticks with the inclusion of an elastic bending (discrete WLC). This discrete model improves the more naive freely jointed chain (FJC) model [3], composed by rigid sticks connected each other that can freely rotate, i.e. that do not include any bending potential. Behind its simplicity, the importance of the FJC model is very high. In fact many flexible polymeric structures can be modeled as FJC as their resistance to bend is very low. An example is the single stranded DNA (ssDNA) whose characteristic elongation as a function of the stretching force can be satisfactorily described as a polymer without bending potential [1, 4]. To take into account the longitudinal elasticity of the polymers, the FJC – as well as the WLC model – needs a correction term not included in its simpler form. This phenomenological correction has been introduced by Odijk [5] in a WLC model by replacing the sticks with harmonic springs, and consists in adding the simple elastic contribution *f*/(*kl*_0_) to the statistical end-to-end distance of the chain with *inextensible* bonds, with *f* the applied force, *k* the elastic constant, and *l*_0_ the Kuhn length of the polymer. In this rigid case, the exact FJC end-to-end distance of the polymer, normalized with its contour length, is the Langevin function 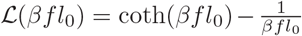. So, the *extensible* FJC (EFJC) presents a normalized end-to-end distance:

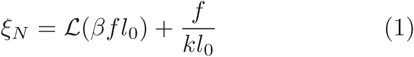

where *β* = 1/*k*_B_*T*, with *k*_B_ the Boltzmann constant and *T* the temperature of the system. This expression has been recently used in fitting the experimental elasticity properties of some polymers [6, 7].

In the same direction, a different choice has also been used by adding the same *f*/(*kl*_0_) term multiplied by a factor given by the very Langevin function:

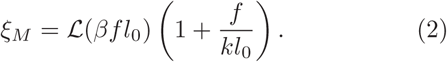

This expression, the first used to fit the experimental outcomes of the Bustamante group, has been, and still is, extensively used to fit the data of different ssDNA and polymer chains [4, 8–16]

Equations (1) and (2) are attractively simple and handy, and they are, in fact, reference formulas for extensible FJC polymer models.

However, they present two big problems. On the one hand – and strangely enough – there is not in the literature a satisfactory derivation of the above formulas from statistical mechanics principles, nor even at high force limits, and this fact even justifies the presence of the two expressions that may adapt to different needs. Some works have presented a formal setting up of the FJC statistical mechanics model, but none has presented a closed exact derivation [17, 18].

On the other hand, though the two expressions appear at a first sight to reasonably agree with the experimental data, that agreement strongly depends on the value of the elastic constant of the polymer studied, which is the main parameter to fit. Generally, the fit analysis of the data by using Eq. (1) are quite imprecise in modeling an EFJC model, and even worse with the use of Eq. (2).

This paper faces both the aforementioned problems by presenting a rigorous analytical derivation of the partition function of the extensible FJC model and, consequently, the exact end-to-end distance *ξ* as a function of the force at all *k* values. An approximated formulas valid only at high forces has also been deduced by means of a complementary derivation.

Then, the analytical findings have been compared with the computer simulations of the Langevin dynamics of an EFJC polymer that moves in a fluctuating environment, reporting an excellent agreement between simulations and the expressions proposed.

## The model

The Hamiltonian of the system is then:

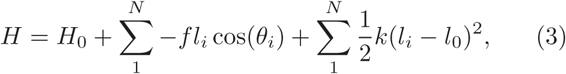

with *N* the number of links, and *l*_0_ the rest length of the spring, which corresponds to the Kuhn length of the polymer. 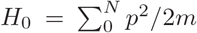 is the kinetic energy contribution. The partition function is then the sum over all the polymer configurations of *e*^−*βH*^, specifically the spatial angles and spring length:

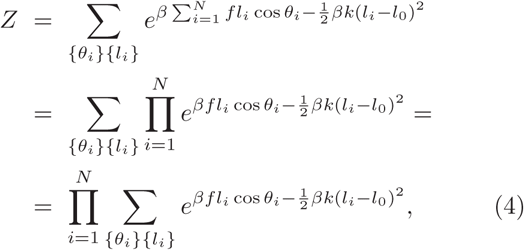

where the kinetic energy contributes with a force independent multiplicative term, here omitted because it is not influent. All the angle configurations are independent from each other, then the partition function is factorized in the *N* equal terms of the above product. So:

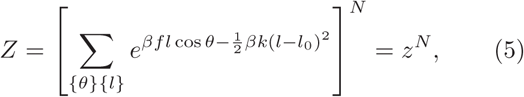

where *z* is the partition function of just one segment.

Given the continuous nature of both the angle values and the spring length, the above expression can be calculated as a spatial integral with volume element *d*Ω = *l*^2^ sin *θ dldθdϕ*:

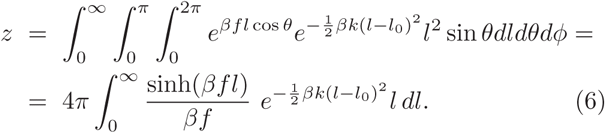

With the change of variable *β fl* = *x*, the integral of Eq. (6) can be rewritten as

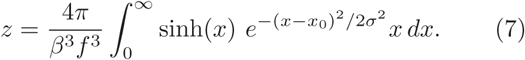

with *σ*^2^ = *β f*^2^/*k*. This integral can be calculated by explicitly writing down the hyperbolic sine and making use of the tabulated integral 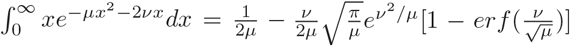. Unfortunately, the presence of the error function *er f*(·) makes the formal outcome useless for practical purposes, yet this expression can be evaluated numerically [17].

## Exact analytical derivation

Nevertheless the above difficulty, the integral of Eq. (7) can be evaluated with a different approach, by writing it as

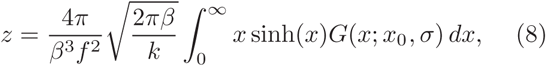

where the Gaussian term 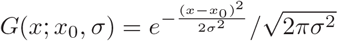 can be expanded in a series of *δ*-functions:

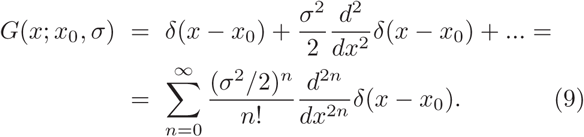

The integral can then be formally written (Weiestrass transform) as

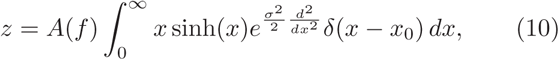

with 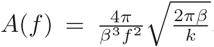. Because of the properties of the *δ*-function inside the integral, *i.e.* 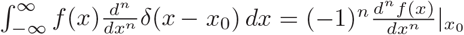 the resulting expression is:

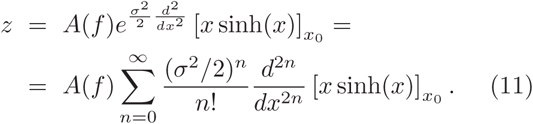

The above expansion is an exact closed form for the partition function.

The general term of the above derivative is

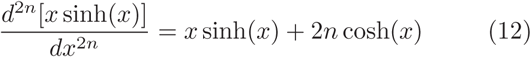

from which, summing up all the terms, we finally obtain:

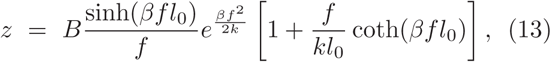

with 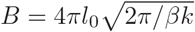. This is the exact partition function of the extensible FJC model.

## High force approximation

For high forces, it is possible to obtain an independent approximation of the partition function. At that limit, the hyperbolic sine of the integral in Eq. (6) can be substituted by the exponential with the positive exponent only:

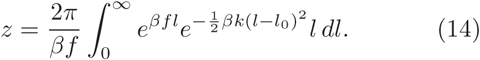

With the variable change *y* = *l* − *l*_0_ − *f*/*k* the integral becomes:

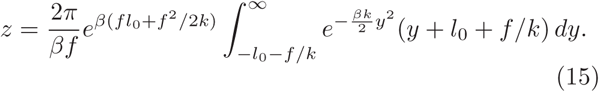

As the force *f* increases, the lower extreme of the integral shifts toward lower values, so permitting a straightforward approximation to –∞ because of the sharpness of the Gaussian integrand. Together with it, the odd term in the integral vanishes, obtaining the simple expression:

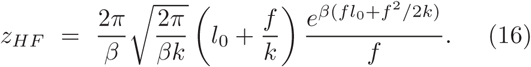

## End-to-end distance

The normalized end-to-end distance along the direction of the force is given by the average:

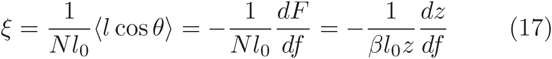

where *F* is the Helmholtz free energy *F* = −1/*β* log *Z*.

With this expression, the value of *ξ*, by using the partition function above evaluated, reads:

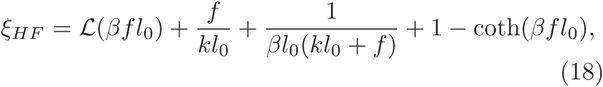

for high forces.

Similarly, the *exact analytical expression* can be calculated by using Eq. (13), obtaining

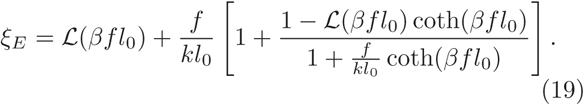

This latter formula is the main result of this paper.

## Langevin simulations

In order to check the analytical result of equation (19) we have performed some dynamical computer simulation. In accordance with the FJC model, the polymer simulated consists of *N* + 1 dimensionless monomers connected by harmonic springs:

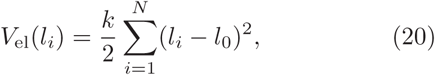

where *k* is the elastic constant, *l_i_* = |**l**_*i*_| = |**r**_*i*+1_ – **r**_*i*_|, is the distance between the monomer *i* + 1 and *i*, with **r**_*i*_ the position of the *i*-th particle, and *l*_0_ is the equilibrium distance between adjacent monomers.

The dynamics of the chain is given by the overdamped Langevin equation of motion

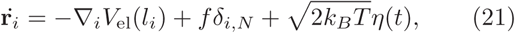

where *η*(*t*) represents the thermal contribution as a Gaussian uncorrelated noise: 〈*η*(*t*)〉 = 0, and 〈*η*(*t*)*η*(*t′*)〉 = *δ*(*t*−*t′*). The nabla operator is defined as ▽_*i*_ = ∂/∂*x_i_***i** + ∂/∂*y_i_***j** + ∂/∂*z_i_***k**. The constant force *f* pulls the last monomer in order to stretch dynamically the polymer, while the first monomer is held fixed.

Fig. 1 shows the extension *ξ vs* the dimensionless applied force *f̃* = *βfl*_0_, obtained from the simulations (symbols), and from the analytical formulas of Eq. (1), Eq. (2), Eq. (18), and Eq. (19) (lines), for different values of the elastic constant parameter 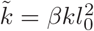. As visible in there, the numerical evaluation of the exact expression *ξ_E_* completely reproduces the simulation data at all the curve extensions. Similarly, the approximation *ξ_H_* correctly approaches the curve at high forces. The figure also shows that the naive approximation *ξ_N_*, while it reproduces the general behavior, lies constantly below the exact expression. The other phenomenological curve *ξ_M_*, lies even lower than the previous one. These differences, very well visible for *k̃* = 3 and *k̃* = 10, remain – though not clearly visible in the plot – as the elastic constant *k̃* increases. The inset of figure 1 reports the differences between the exact *ξ_E_* and the two phenomenological expressions *ξ_N_* and *ξ_M_*, as a function of *f̃*, for *k̃* = 10. We can notice there that the difference Δ*ξ_N_* tends to zero for *k̃* → ∞, while Δ*ξ_M_* tends to the value 1/ *k̃*. The difference between the curves is also present at low forces, where the three lineal approximations read: 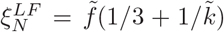, 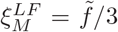, and 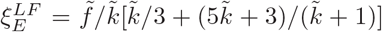, revealing very different slope behaviors. The expression 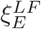 can be used for fit purposes by using low force data.

**FIG. 1:**
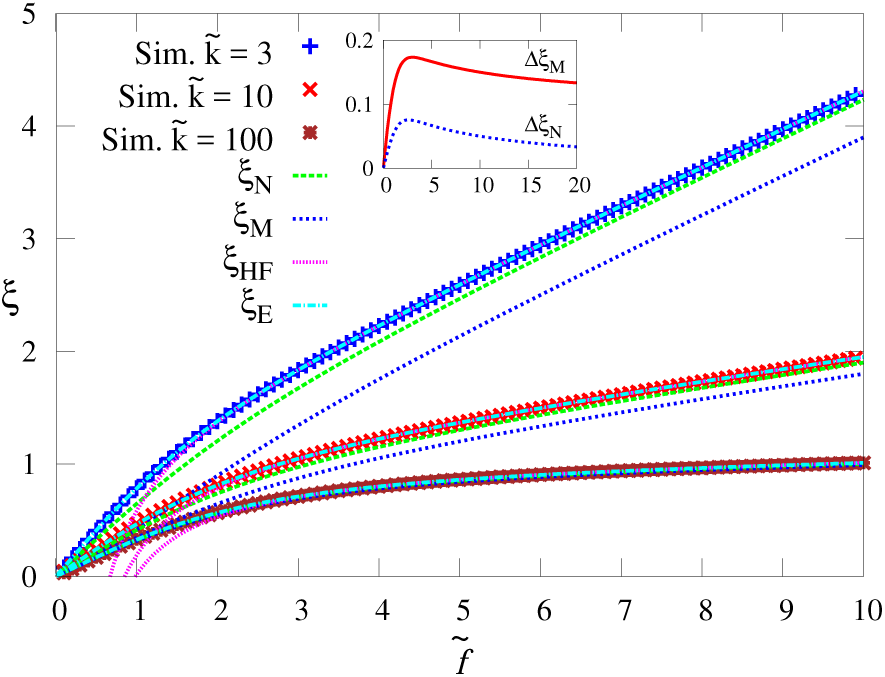
Normalized extension *ξ* as a function of the dimensionless force *f̃* = *βfl*_0_ for three values of the dimensionless elastic constant 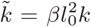 in the extensible FJC model. The symbols represent the data from the simulations, and the lines the analytical expressions defined in the text. Inset: the difference between Eq. (19) and Eq. (2) (Δ*ξ_M_*), and between Eq. (19) and Eq. (1) (Δ*ξ_M_*), with *k̃* = 10.

In order to estimate the effectiveness of the formulas obtained, we have performed a numerical fit on the simulations for different values of *k̃* considered as a free parameter. The results are shown in Table I. As evident there, the predicted *k̃*s present an error up to 22% for Eq. (1), and even up to 32% for Eq. (2). Such big errors could be the origin of some anomalous discrepancies in the estimations of *k* [19]. The same fit analysis has also been done by using an ensemble of data points obtained from the numerical estimation of the end-to-end distance obtained by the exact integral (Eq. (6)). The results of the fitted parameters *k̃* for the phenomenological expressions (Eq. (1) and Eq. (2)) gives errors comparable with those presented in Table (I), with a difference of a few percents. Only for *k̃* = 1000 the error decreases of around 40%. The exact formula reports, expectably, negligible errors in the predicted *k̃* values.

**TABLE I:**
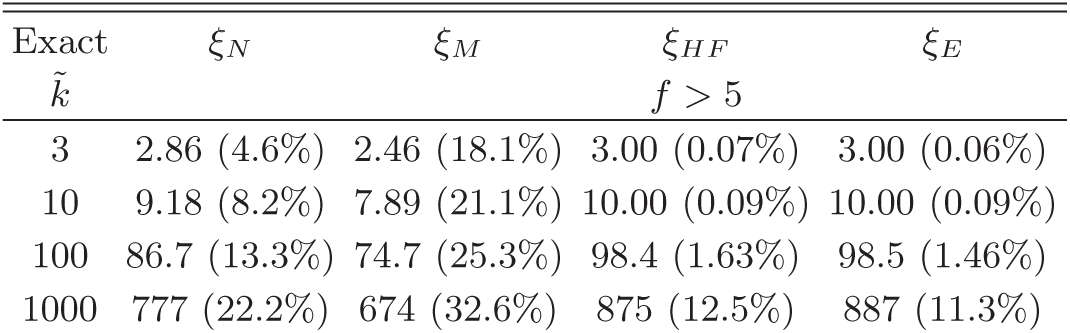
Values of *k̃* obtained by fitting the simulations data for the real parameter value listed in the first column of the table. In parenthesis, the error with respect the exact value.

## Discussion

This paper presents the analytical derivation from statistical mechanics principle of the partition function of the extensible FJC model from which the exact end-to-end distance as a function of the stretch force has been derived. An approximated formula valid at high forces only has also been presented by means of a complementary derivation, and a double check by means of Langevin simulations has been performed.

The expression here derived establishes the EFJC as the most complex analytically-solvable polymer model.

The formula obtained is a combination of elementary functions simple enough to be implemented in any fit of experimental data of flexible polymers.

## Acknowledgments

This work is supported by the Spanish projects Mineco No. FIS2017-87519-P and No. FIS2014-55867-P, both co-financed by FEDER funds. We also thank the support of the Aragón Government and Fondo Social Europeo to FENOL group. We also want to thank Mario Floría for useful discussions.

